# Another look at microbe–metabolite interactions: how scale invariant correlations can outperform a neural network

**DOI:** 10.1101/847475

**Authors:** Thomas P. Quinn, Ionas Erb

## Abstract

Many scientists are now interested in studying the correlative relationships between microbes and metabolites. However, these kinds of analyses are complicated by the compositional (i.e., relative) nature of the data. Recently, Morton et al. proposed a neural network architecture called mmvec to predict metabolite abundances from microbe presence. They introduce this method as a scale invariant solution to the integration of multi-omics compositional data, and claim that “mmvec is the only method robust to scale deviations”. We do not doubt the utility of mmvec, but write in defense of simple linear statistics. In fact, when used correctly, correlation and proportionality can actually outperform the mmvec neural network.

## 1 Response

Many scientists are now interested in studying the correlative relationships between microbes and metabolites (e.g., [1, 2, 3, 4]). However, these kinds of analyses are complicated by the compositional (i.e., relative) nature of the data [5, 6]. Recently, Morton et al. proposed a neural network architecture called mmvec to predict metabolite abundances from microbe presence [7]. They introduce this method as a scale invariant solution to the integration of multi-omics compositional data, and claim that “mmvec is the only method robust to scale deviations”. We do not doubt the utility of mmvec, but write in defense of simple linear statistics. In fact, when used correctly, correlation and proportionality can actually outperform the mmvec neural network.

Scale invariance is important because we do not want a method that is sensitive to (i.e., is variant to) changes in technical factors like sequencing depth (i.e., differences in scale). In compositional data analysis (CoDA), scale invariance is forced by using a log-ratio transformation that recasts the data with respect to an internal reference [8]. The resultant log-ratios are scale invariant, and so any analysis of log-ratios is scale invariant. This is true for multi-omics data too, but only if the transformation is performed correctly. Let us consider two possible transformations of the multi-omics data, presented here as functions of the input:

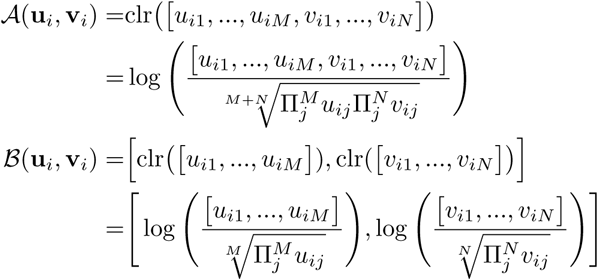

for sample *i*, where **u**_*i*_ measures the 1…*M* microbes and **v**_*i*_ measures the 1…*N* metabolites. Only approach 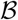 is scale invariant, but Morton et al. use approach 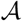 when they claim that correlation and proportionality are unreliable.

Why is approach 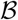 valid, but not approach 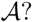 It is because the microbe and metabolite data are generated from two separate sampling processes: they are individually, not jointly, constrained to sum to 1. In other words, the abundance of microbe 1 is limited by the abundance of microbes 2-to-*M*, but is in no way limited by the abundance of metabolites 1-to-*N*. Consequently, the denominator from approach 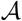 has no meaning. On the other hand, the denominators from approach have the property that they cancel any constant factor multiplied with the sample values in each numerator. As such, they cancel the implicit biases that arise from the sequencing procedure and cause the samples to be on different scales. An additional property of these numerators is that they are useful normalization factors themselves [9]: under the assumption that the majority of features are unchanged, approach 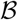 will make the transformed data proportional to the original (absolute) data (thus performing an effective library-size normalization).

How important is the choice between approach 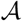 and approach 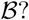 We repeated the authors’ analysis to measure the F1-score (precision and recall) for the top microbe-metabolite associations, except this time we used approach 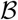. Figure 1 shows the updated performance of correlation and proportionality, both of which outperform mmvec on their simulated benchmark. Interestingly, correlation (Spearman and Pearson) performed best, suggesting that the “ground truth” includes power-law relationships between microbes and metabolites (i.e., log-linear relationships with slopes other than 1). Since *ϕ* and *ρ* are designed for intercept-free linear relationships, pairs in which one feature is proportional to another when taken to an exponent will usually go undetected.

**Figure 1:**
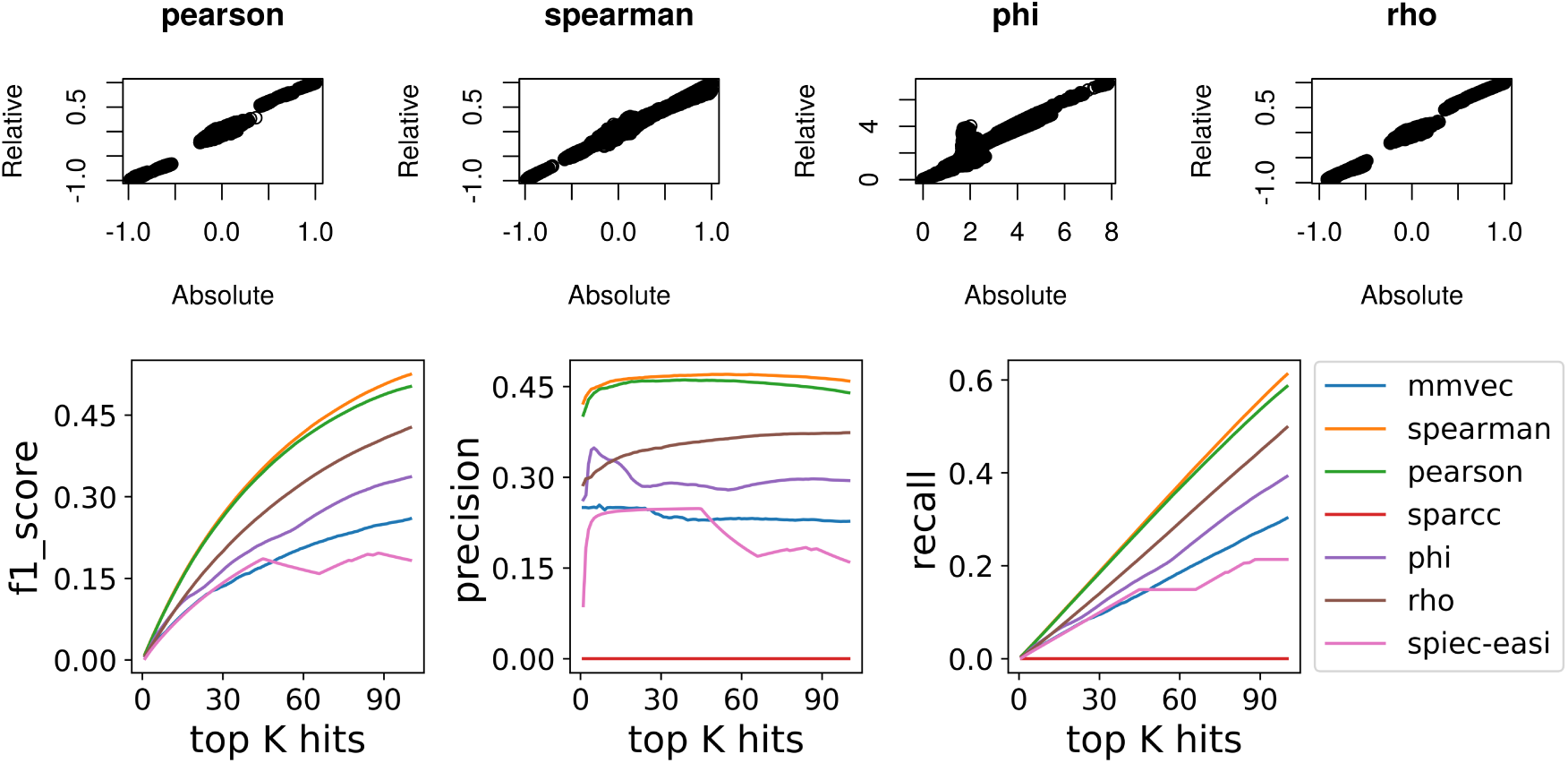
Our re-analysis of the Morton et al. simulated data shows that, with the correct log-ratio transformation, simple linear statistics are scale invariant and sufficient for use in the integration of multi-omics compositional data. In the top panels, we see excellent agreement between absolute and relative correlation, as well as between absolute and relative proportionality. In the bottom panels, we see the updated performances from the simulated data benchmark. When used correctly, correlation and proportionality can actually outperform the mmvec neural network. All scripts available from https://doi.org/10.5281/zenodo.3544999.

We do not disagree that neural networks can add value to multi-omics data integration. Their ability to learn non-linear relationships could improve metabolite prediction by directly modeling complex microbe-microbe interactions. However, neural networks do not offer a magical solution to the problems of compositional data analysis [10]. They are merely a nested series of transformed linear operators. As such, they may be prone to yield spurious results whenever a simple linear method would. It seems to us that mmvec’s primary advantage is how it handles the compositional data, not its neural network architecture. Indeed, our analysis shows that when we transform the multi-omics data correctly, simple linear methods outperform mmvec in their own benchmark. We conclude by reminding our readers that not all problems in biology are solved by adding layers of complexity: sometimes it is sufficient to use our simplest solutions more carefully.

